# Light-evoked activity and BDNF regulate mitochondrial dynamics and mitochondrial localized translation

**DOI:** 10.1101/438739

**Authors:** Alex Kreymerman, Jessica E. Weinstein, Sahil H. Shah, David N. Buickians, Anne Faust, Yolandi Van Der Merwe, Michael M. Nahmou, In-Jae Cho, Star K. Huynh, Sonya Verma, Xiao-Lu Xin, Michael B. Steketee, Jeffrey L. Goldberg

**Affiliations:** Byers Eye Institute, Stanford University, Palo Alto, CA 94303; University of Miami Miller School of Medicine, Miami, FL 33136; Medical Scientist Training Program, University of California, San Diego, CA 92093; Department of Ophthalmology and McGowan Institute for Regenerative Medicine, University of Pittsburgh, Pittsburgh, PA 15213

## Abstract

Mitochondria coordinate diverse functions within neurites, including signaling events for axonal maintenance, and degeneration. However, less is known about the role of mitochondria in axon development and maturation. Here we find that in maturing retinal ganglion cells (RGCs) in vivo, axonal mitochondria increase in size, number, and total area throughout development. We demonstrate through multiple approaches in vivo that the mechanism underlying these mitochondrial changes are dependent on eye opening and associated neuronal activity, which can be mimicked by brain derived neurotrophic factor (BDNF). We report downstream gene and protein expression changes consistent with mitochondrial biogenesis and energetics pathways, and present evidence that the associated transcripts are localized and translated at mitochondria within axons in an activity-dependent manner. Together these data support a novel model for mitochondrial-localized translation in support of intra-axonal mitochondrial dynamics and axonal maturation.

## Introduction

Neurons are among the highest metabolically active cell types in the body. This is due in part to mitochondrial oxidative phosphorylation highly coupled with the energy demand generated by electrophysiologic activity and associated signaling^1-3^. Beyond ATP production, mitochondrial activities such as calcium homeostasis, fatty acid oxidation, secondary messenger and signaling pathway modulation also participate in supporting neurons and their extensive axonal compartments^4^. Mitochondrial activities are critically dependent on the expression and assembly of approximately 600-1500 proteins encoded in the nucleus^5-12^, yet mitochondria can be separated down the axon by a meter or more from the cell body^13^.

As a result, such cells have evolved unique mechanisms for maintaining continuous communication between mitochondria and the nucleus^14^. These include shuttling mitochondria and their nuclear-encoded proteins up and down axons using motor proteins kinesins and dyneins^15-18^. Transported mitochondria are also capable of undergoing fusion or fission with neighboring mitochondria, acquiring or shedding genetic material and proteins^19^. Finally, new mitochondria can also be assembled and packaged with nuclear and mitochondrial encoded proteins, in a process known as mitochondrial biogenesis. This process takes place in the perinuclear space and within axons, leading to increased numbers of mitochondria in neuronal compartments^20-22^. Together these changes in mitochondrial localization, size and total cellular volume are referred to as mitochondrial dynamics.

An additional mechanism implicated in supplying nuclear proteins to distal axonal mitochondria is the transport and then local translation of RNA in axonal compartments (reviewed elsewhere^23^). Interestingly, many investigations indicate that a consistent and major portion of axon-localized transcripts encode nuclear proteins that regulate mitochondrial functions^24-27^. Additionally, nuclear-encoded mitochondrial transcripts have been shown to physically localize on/in mitochondrial membranes^28-32^, with further evidence suggesting that mitochondria can act as local translation sites^33-35^. However, it is not yet known how such localization is regulated. Here we find that developmental changes in axonal mitochondria are regulated by activity in vivo, and explore associated regulation of mRNA expression and localization by activity in RGCs in vitro.

## Results

### Mitochondrial networks reorganize at the time of eye opening

We first studied mitochondrial organization in RGC axons in transgenic mice expressing cyan fluorescent protein (CFP) fused to the COX8a mitochondrial targeting sequence under control of the Thy-1 promoter (Thy1-CFP/COX8A). In this mouse, approximately 5% of RGCs express the CFP/COX8a transgene, permitting visualization and quantification of mitochondria in RGC axons (Figure 1A,B). We used this mouse model to investigate axon-specific mitochondrial networks at postnatal (P) days 9, 12, 15, and 45, as RGCs experience significant developmental changes through this time period^28-31,36-39,40-44^. Of note, these time points also follow the period of developmental cell death in RGCs, which peaks at P5 in mice ^45-47^, thus allowing for the identification of mitochondrial changes independent of cell death signaling, which can influence mitochondrial morphology^48,49^. Analysis of CFP-labeled mitochondria in whole mount retinas and optic nerves by confocal microscopy revealed significant reorganization in RGC axons throughout postnatal development (Figure 1B). Overall, mitochondria increased in size, number, and occupied a greater percentage of axonal area from P9 to P45 (Figure 1C-E). Interestingly, within a relatively short window of development, around eye opening (P12/13 to P15), mitochondrial size, number, and occupied area increased in both RGC retinal and optic nerve axon segments, with optic nerve mitochondria experiencing the greatest change during this time window.

**Figure 1.**
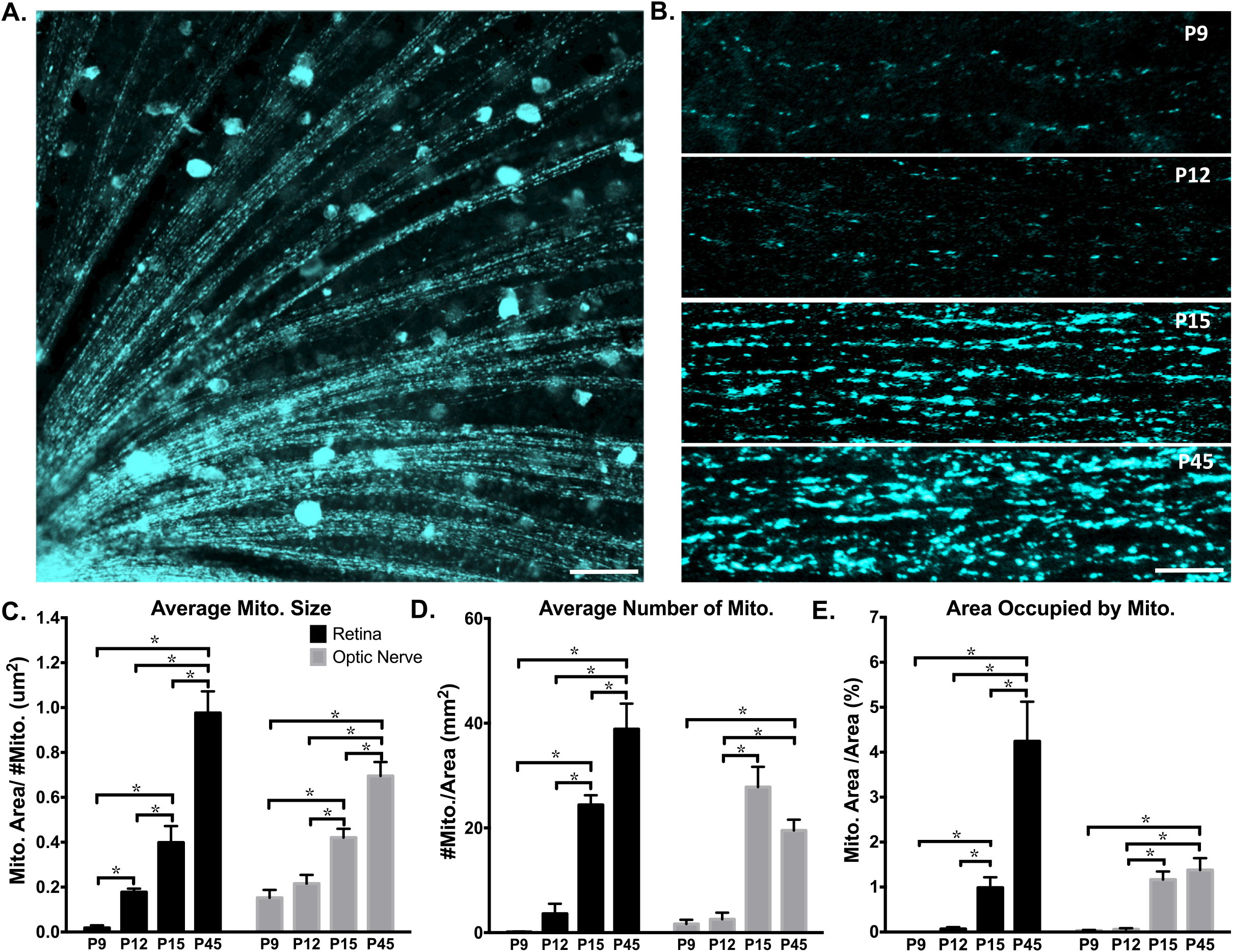
Axonal mitochondria increase in size, number and area through eye development. CFP+ mitochondria were imaged by confocal microscopy and analyzed in ImageJ. (A) Example image of mitochondrial labeling within retinal RGC axon segments (50um scale bar) and (B) in optic nerve RGC axon segments from postnatal day 9 (P9), P12, P15, and P45 mice (lum scale bar). (C-E) In both retinal and optic nerve RGC axons, the average mitochondrial size, number, and area (measured as percent of cross sections, representing fractional volume) increased from P9 to adulthood. (Error bars indicate SEM; N≥ 3 mice per age, with 9 images analyzed per animal; one-way ANOVA with Holm-Sidak correction for multiple comparisons, * p ≤ 0.05.)

Although Thy-1 gene expression peaks around P12 in RGCs and continues to be stably expressed throughout adulthood^50^, and thus Thy-1 promoter-related artifacts are unlikely, mitochondria size changes from P12-P15 were also confirmed by transmission electron microscopy (TEM; Figure 2). Specifically, mitochondrial size, number, and occupied area increased in RGC axons from P12 to P15 by 25%, 105%, and 31% respectively (Figure 2B, C, D). Thus by both fluorescence and TEM imaging, RGC axon mitochondrial size, number, and occupied area increase during this developmental window.

**Figure 2.**
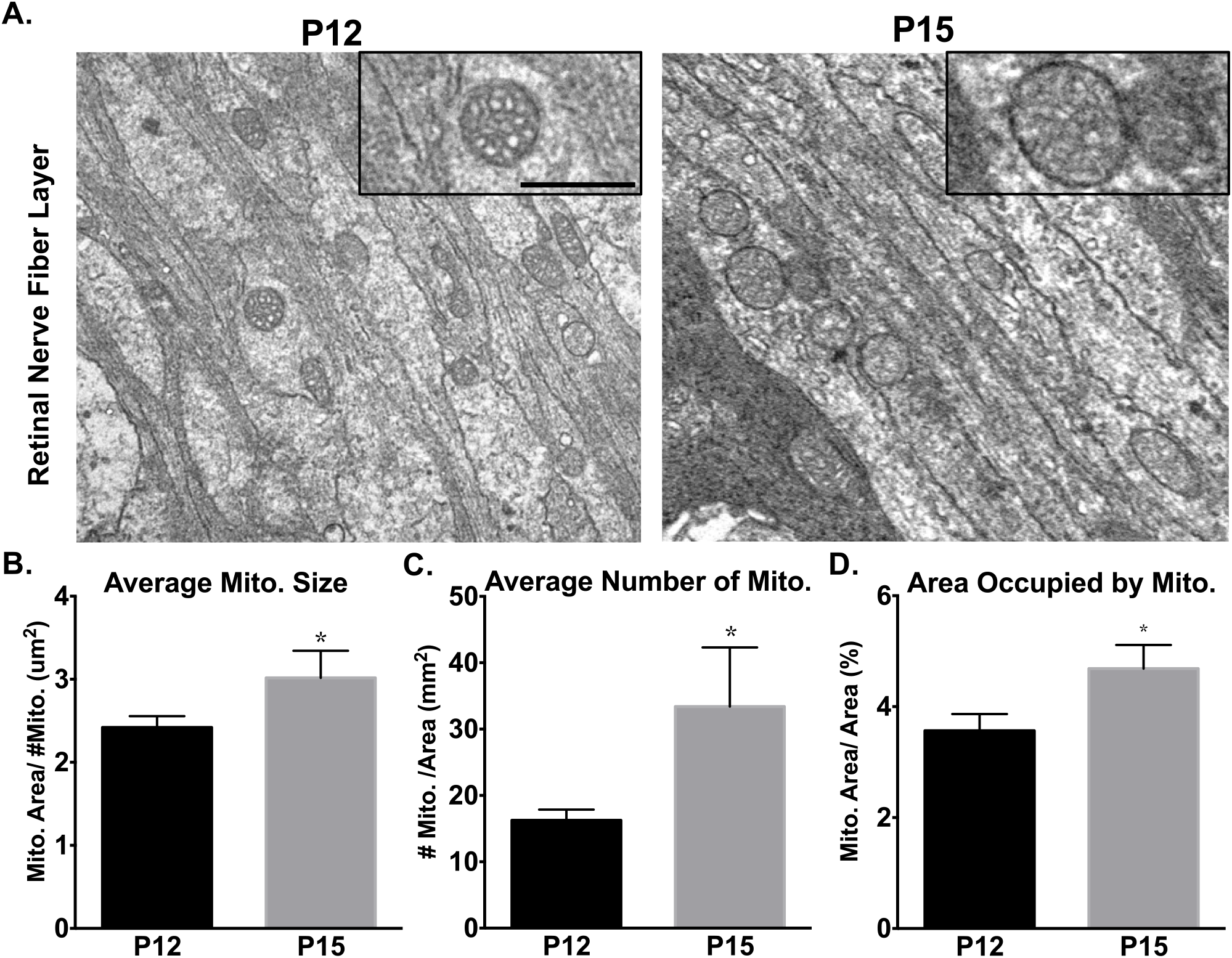
Mitochondrial size and number increase at eye opening in RGC axons. (A) Mitochondria were imaged and quantified in RGC optic nerve axons by transmission electron microscopy before (P12) and after (P15) eye opening. Increased magnification (insets) shows mitochondrial membrane, cristae, and representative mitochondrial size differences. Scale bar 500 nm. (B) Average mitochondrial size, (C) number and (D) area increased significantly between P12 and P15. (Error bars indicate SEM; n ≥ 30 sections; t-test * p < 0.05).

### Eye opening regulates mitochondrial networks

As eye opening occurs between P12-P15 with a concomitant significant increase in visual activity, we asked whether the mitochondrial morphological changes are dependent on eye opening in CFP-COX8a mice with surgically premature or delayed eye opening. (Figure 3A). For premature eye opening, we surgically opened the P10 eyelid margin, two days prior to normal eye opening, and allowed animals to mature to P12. We found increases in mitochondrial size, number and occupied area in retinal and optic nerve axons compared to unopened P12 eyes (Figure 3B, C, D). Thus eye opening accelerates the mitochondrial morphology changes identified in normal development.

**Figure 3.**
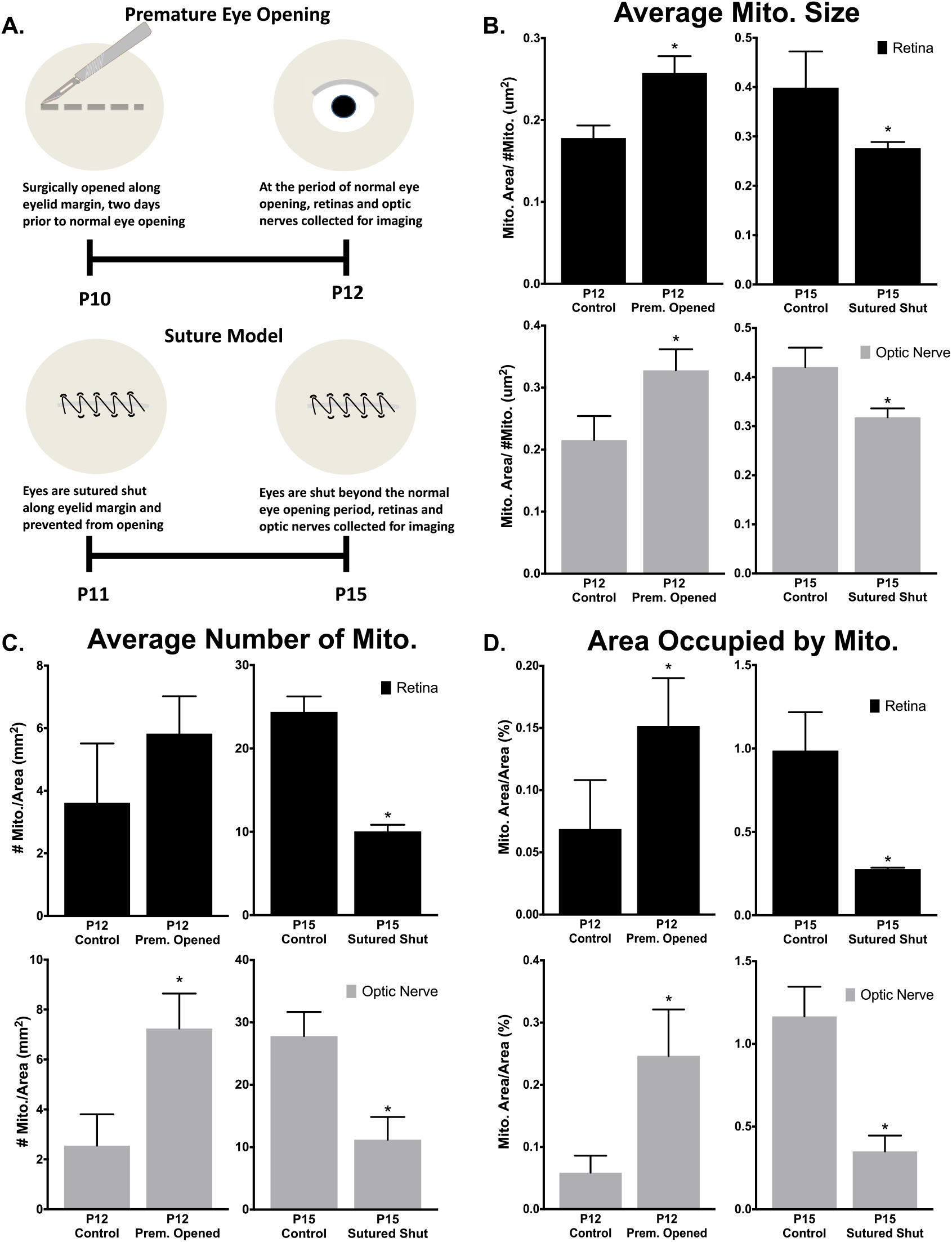
Eye opening is sufficient and necessary developmental changes in mitochondrial size and localization. (A) Surgical model for premature eye opening and sutured eyelid closure. (B) Average mitochondrial size, (C) number, and (D) area increase with premature eye opening, and this developmental increase is inhibited by prolonged eye closure. (Error bars indicate SEM; N≥ 3 mice per condition, 9 images analyzed per animal; Students t-test * p < 0.05.)

Conversely, to determine if eye opening is necessary for the developmental increases in mitochondrial size, number, and occupied area from P12-15, we delayed eye opening by suturing eyelids shut at P11, prior to natural eye opening, then allowed mice to mature to P15, delaying eye opening by 2-3 days (Figure 3A). Compared to age-matched controls, the delayed eye opening model led to significant decreases in all mitochondrial measurements (Figure 3B, C, D), in most cases back to levels found in P12 animals. Thus, these data suggest that the process of eye opening is sufficient and necessary for the mitochondrial network increases found from P12-15.

### RGC activity and BDNF regulate mitochondrial networks

These findings suggested the hypothesis that light-stimulated electrical activity in axons (i.e., action potentials) contribute to the observed changes in axon mitochondrial distribution and morphology. To test the contribution of electrical activity to axonal mitochondrial changes during the period of eye opening, we pharmacologically inhibited both spontaneous and light-evoked electrical activity in RGCs by intravitreally injecting tetrodotoxin (TTX)^51^ prior to eye opening at P11 and again after eye opening at P13, followed by mitochondrial quantification at P15. We found that TTX but not control vehicle injection inhibited mitochondrial increases in size in retinal and optic nerve axons (Figure 4A, B), and mitochondrial number and occupied area only in the optic nerve portion of RGC axons (Figure 4C, D), in all cases to levels equivalent to vehicle injected P12 animals. Thus RGC electrical activity is a significant contributor to mitochondrial network changes that occur concomitant with eye opening. However, discrepancies in mitochondrial numbers and area within retinal versus optic nerve axons likely indicate additional regulation that contributes to mitochondrial dynamics during this period.

**Figure 4.**
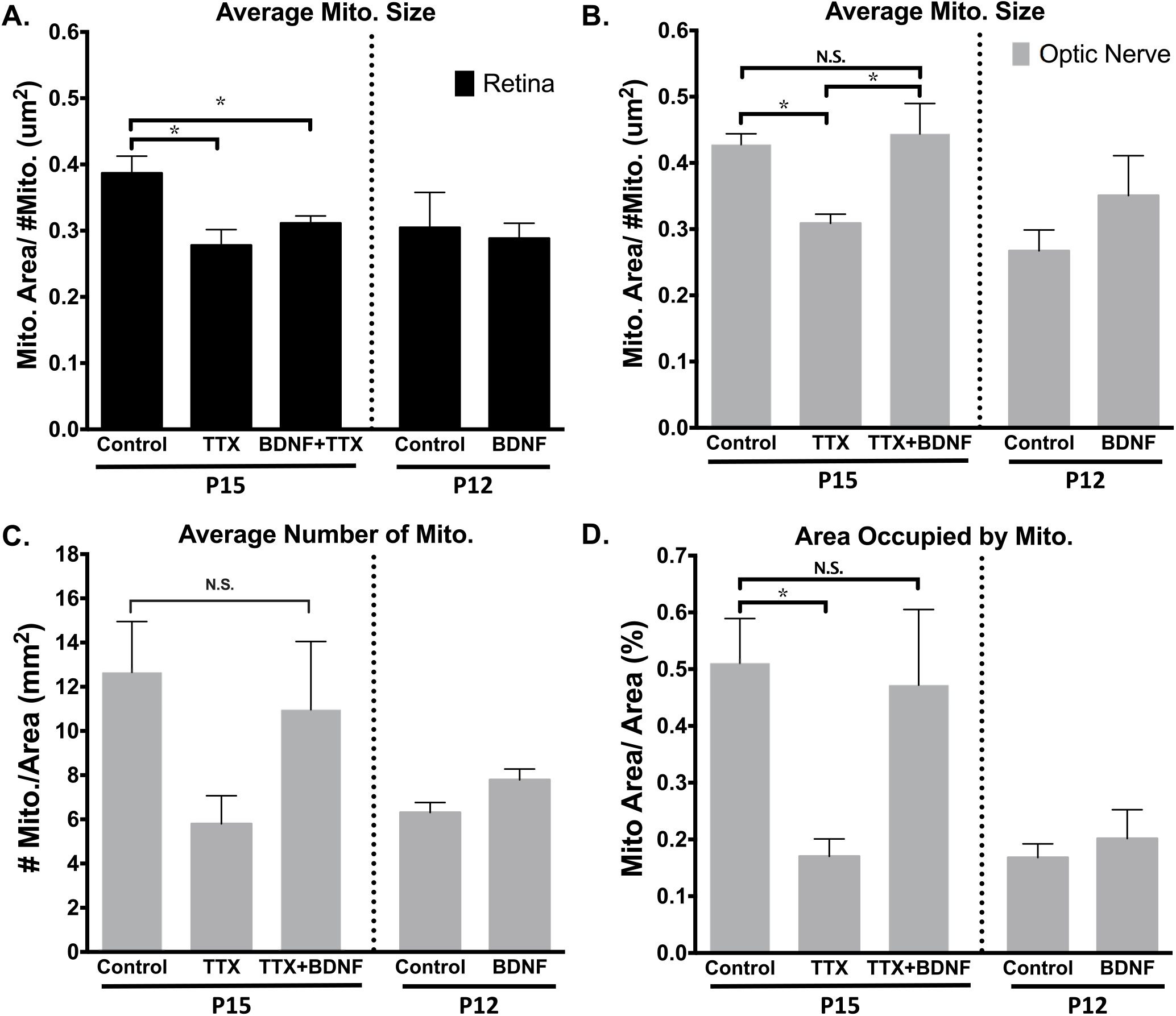
Mitochondrial developmental changes are dependent on retinal electrical activity and are partially rescued by BDNF in optic nerve axons. Control, TTX- or TTX plus BDNF-treated mice analyzed at P15, as well as control or BDNF-treated mice analyzed at P12 are graphed on the same axis for comparison, but the experiments were performed and analyzed separately. Measured changes in (A) average mitochondrial size within retinal axons and (B) optic nerve axons, as well as the corresponding mitochondrial (C) number and (D) area. (Error bars indicate SEM; N= 3 mice per condition, 9 images analyzed per animal; one-way ANOVA with Newman-Keuls multiple comparisons test (A-C) or Fisher LSD test (D), * p < 0.05)

Downstream of electrical activity, BDNF expression has been shown to be regulated by eye opening and to be blocked by TTX injection^36^, and to modulate mitochondrial dynamics^36, 52-55^. To determine whether BDNF could rescue the inhibitory effects of TTX on mitochondrial networks, BDNF and TTX were co-injected at P11 and again at P13, and mitochondrial parameters were measured at P15. BDNF was capable of significantly reversing the TTX-induced decreases in mitochondrial size, and showed a non-significant trend towards such rescue in mitochondrial number and occupied area in optic nerve axons (Figure 4B-D). Of note, this rescue effect of BDNF was only detected in the optic nerve but not retinal axons (Figure 4A), suggesting again that different mechanisms regulate mitochondrial dynamics in a compartment-specific manner within the axon. Furthermore, when BDNF was injected at P10 and mitochondrial parameters were measured at P12, BDNF did not increase mitochondrial size, number or area on its own (Figure 4A-D). Nonetheless, these data suggest that activity and BDNF are both critical in regulating the morphology and distribution of optic nerve axon mitochondria during this stage of visual system development.

### Activity and BDNF regulate the expression of nuclear encoded mitochondrial genes

To investigate molecular mechanisms associated with activity and BDNF, we explored the transcriptional influence of exogenously added BDNF and TTX on nuclear-encoded mitochondrial gene expression in RGCs. To accomplish this, we injected TTX and/or BDNF in combination or alone at P11 and P13, acutely purified RGCs from P15 retinas, and extracted RNA for qRT-PCR gene arrays. We then analyzed expression data and conducted pathway analysis. Major upstream regulators were identified by probing the gene expression sets for targets of known regulators, and then filtering for genes whose expression was concordant with the inhibitory effects of TTX and subsequent rescue with BDNF on mitochondrial morphology and distribution (Figure 5A). The resulting analysis revealed that PGC1-α and RICTOR, master mitochondrial dynamics and energetics modulators, were putative upstream regulators of genes modulated by TTX and/or BDNF (Figure 5B, C). For many mitochondria genes regulated by RICTOR and PGC1-α, TTX and BDNF showed opposing effects on expression. In most cases, the gene expression profile of TTX+BDNF mimicked that of BDNF alone, suggesting BDNF’s effects on gene expression were dominant over effects of TTX and placing BDNF downstream of activity. Furthermore, BDNF increased basal and maximum respiratory capacity in purified RGCs in vitro even in the presence of TTX (Figure 5D), consistent with pathways predicted by these gene expression changes. Ontological analysis of these gene expression datasets suggested opposing functions between BDNF and TTX, with fission/fusion and mitochondrial biogenesis pathways induced by BDNF and suppressed by TTX (Figure 5E, F), consistent with our *in vivo* data.

**Figure 5.**
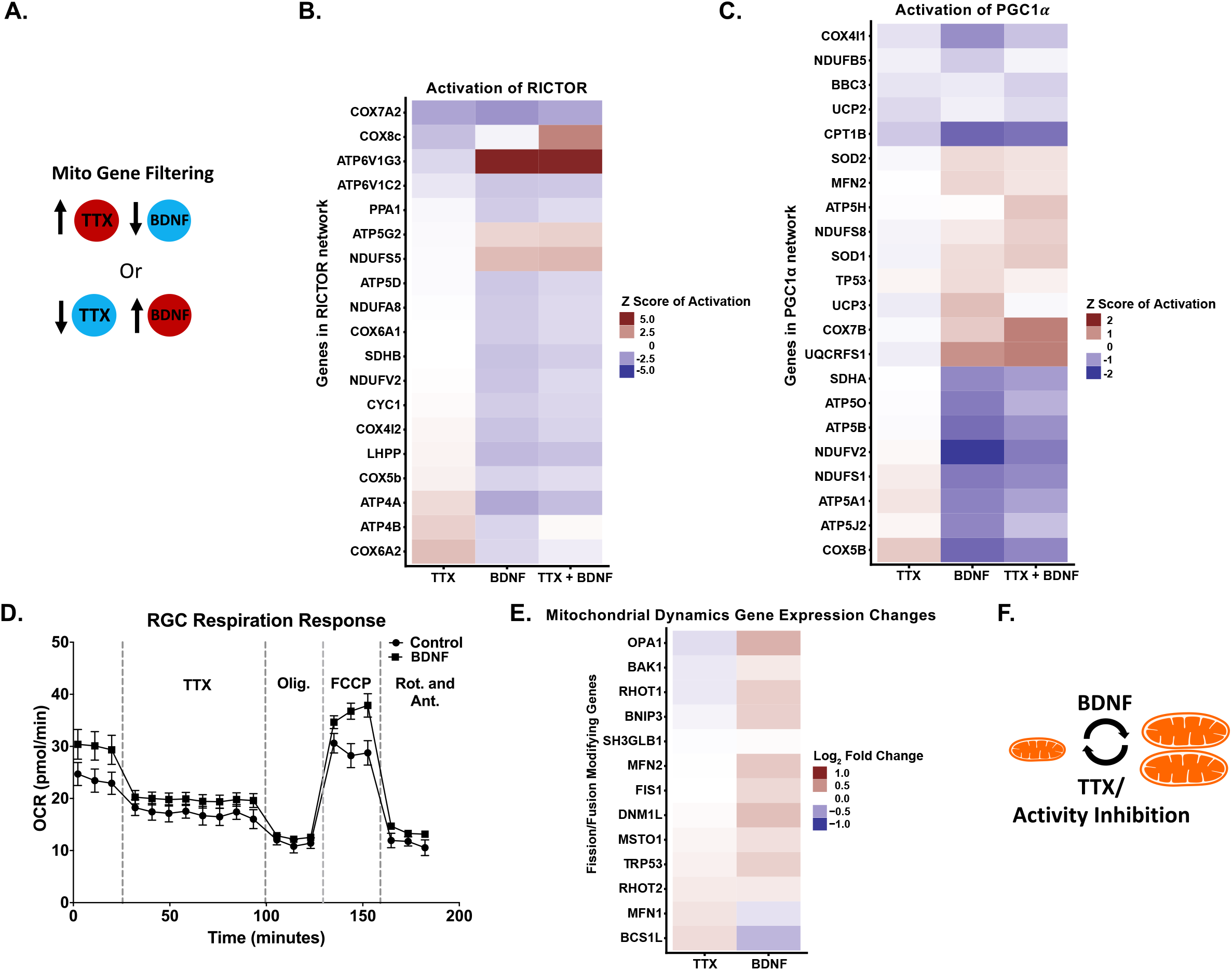
RGC nuclear-encoded mitochondrial gene expression in response to activity inhibition with TTX and/or BDNF is consistent with inhibition versus activation of mitochondrial dynamics and energetics. (A) Model for filtering data acquired from RT-PCR gene array analysis of P15 acutely purified RGCs, after TTX and/or BDNF intravitreal injections at Pll and P13 (N=3 RGC preps per condition). Filters were placed to identify gene expression regulated in opposing directions by BDNF and TTX. The resulting genes were then passed through IP A^®^ pathway analysis software, which suggested 10 major upstream regulators, with PGC1-α and RICTOR at the top of the list. Downstream gene expression data modulated by these upstream regulators were transformed into Z-score of activation. Up- or downregulated gene sets are denoted by color.(B)Genes identified in our array that are regulated by RICTOR represent mainly energetics genes. (C) Genes identified in our array that are regulated by PGCl-α represent mainly mitochondrial dynamics and biogenesis regulators. (D) Measuring the effect of BDNF on mitochondrial dependent oxygen consumption in purified RGCs shows an increase in the basal respiration rate, and maximum respiration capacity (with FCCP addition) regardless of activity inhibition by TTX (introduced 35min after initial recording). TTX, Oligomycin, FCCP, and Rotenone/Antimycin A, were added sequentially at time points marked with vertical lines. Recorded values were acquired using the Seahorse XF96 instrument (Error bars indicate SEM; n=6 replicates per condition, pooled from 3 separate RGC preps and assayed on one plate) (E) Genes identified in our array that have been previously demonstrated as mitochondrial fission/fusion or mitochondrial size modifying are opposingly regulated by TTX and BNDF, with most genes upregulated by BDNF. (F) Model of the predicted mitochondrial events triggered by TTX or BDNF, based on gene pathway analysis and the identified mitochondrial changes in injected mice.

### Activity regulates mitochondrial associated local translation and mitochondrial dynamics

Our gene arrays showed that expression of many of the mitochondrial genes assayed were suppressed by TTX-mediated inhibition of activity. To investigate whether transcription or translation activity were being globally downregulated in RGCs by activity inhibition, we treated RGCs with 5-ethynyluridine (EU), a uridine analog, or O-propargyl-puromycin (OPP), a puromycin analog. These molecules readily incorporate into newly synthesized RNAs (with EU) or proteins (with OPP), and can be conjugated to fluorophores with click chemistry to visualize the location and relative amount of synthesis taking place within cells^56,57^. Using this approach, we first tested whether TTX inhibited transcription or translation by intravitreally injecting TTX at P11 and P13, and then pulsing with EU or OPP at P15 for 1hr. Upon visualization, no detectable differences in signal intensity from EU or OPP were identified in retinas (Figure 6A), suggesting that transcription and translation where not broadly inhibited by activity suppression.

**Figure 6.**
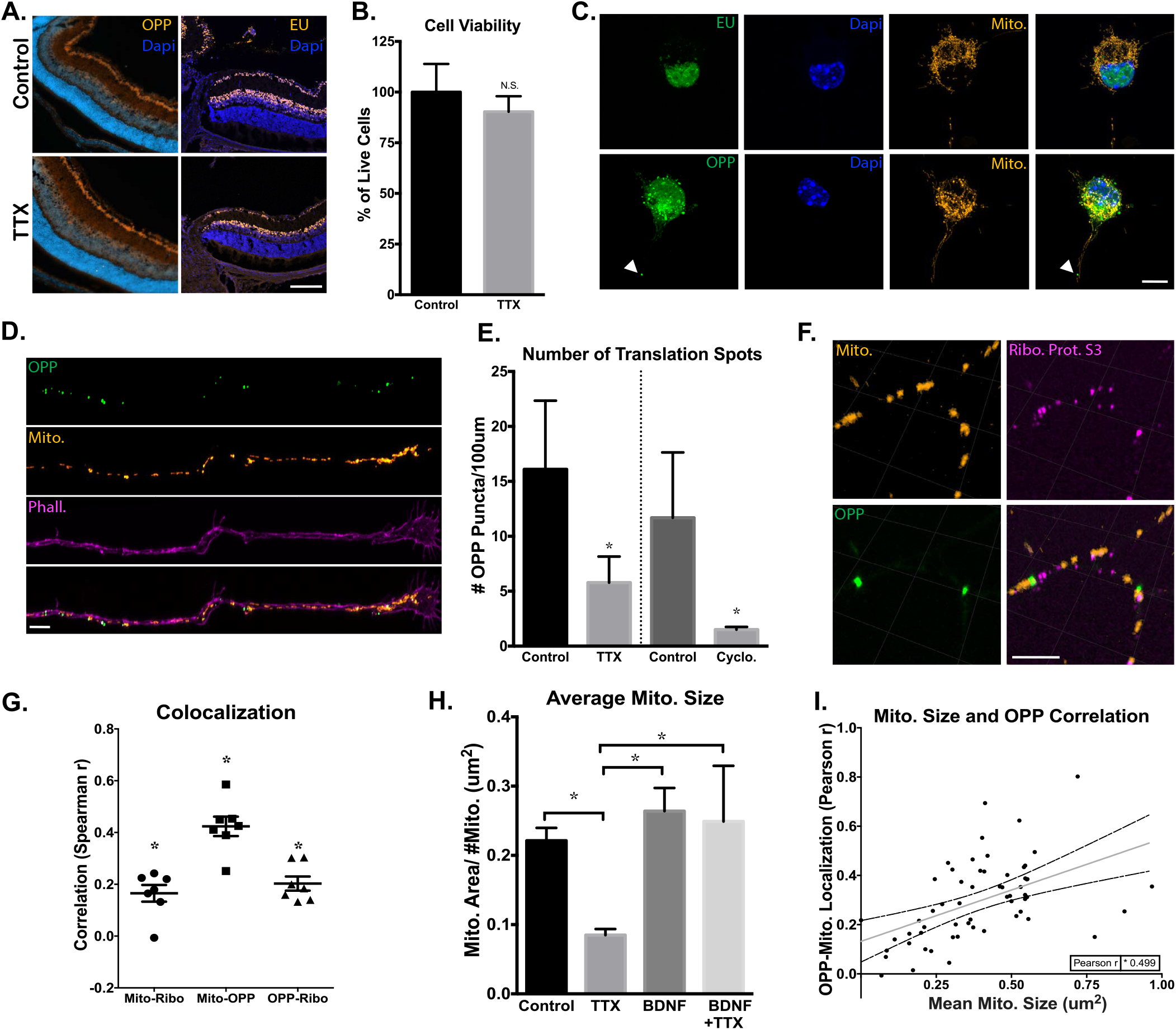
Activity regulates mitochondrial localized protein translation in axons. (A) Representative images collected by confocal microscope of PI 5 retinas, after in vivo intravitreal injections of TTX or BSS (control) at PI 1 and P13, and an injection of EU or OPP 2 hours before dissection and tissue processing. There were no detectable differences in EU or OPP fluorescence in the retina after TTX-mediated inhibition of activity (lOOum scale bar). (B) CeXll viability of TTX- and-control treated RGCs, identified as Calcein-AM positive and Sytox negative, normalized for total cell number by Hoescht, and quantified as percent change relative to control treated cells (Error bars indicate SEM; N=3 replicate RGC preps, n>l 00 cells per replicate condition; Student’s t-test, *p < 0.05.). (C) Confocal images of cultured RGCs treated with BacMam virus, labeling mitochondria with DsRed, and pulsed with EU for lhr or OPP for 15 min before fixation and staining for newly synthesized RNA and protein, respectively. EU RNA and OPP protein staining were strongly detected in RGC nuclei and cell bodies, and newly translated proteins were also detected in axons (marked by arrows, lOum scale bar). (D) Axon tips demonstrate strong OPP+ puncta throughout growth cone and terminal axon domains. OPP-labeled with Alexa488, mitochondria with DsRed, and axon tips with phalloidin Alexa647 (5um scale bar). (E) Quantified average number of OPP puncta per 100 pm of P4 RGC’s axon termini, treated with TTX or cycloheximide and with vehicle treated controls. Groups separated by vertical line were experimentally treated and analyzed separately (Error bars indicate SEM; n>10 randomly imaged axons, selected from 3 replicate RGC preps; Student’s t-test, *p < 0.05). (F) Representative image of mitochondria, OPP-labeled new protein synthesis, and ribosomal protein S3 colocalization within axons (lum scale bar), along with (G) the spearman correlation values for the association of OPP to Mito, Mito to Ribosomes, and Ribosomes to OPP signals in line-scanned axons (Error bars indicate SEM; n=7 randomly imaged axons, selected from 3 replicate RGC preps; individual R values were all significant, * p < 0.05). (H) Quantified mean mitochondrial size in axons from RGCs incubated with TTX, BDNF, TTX and BDNF, or vehicle controls (Error bars indicate SEM; n>10 randomly imaged axons, from 3 replicate RGC preps; one-way ANOVA with Holm-Sidak’s test, *p < 0.05). (1) Pearson’s correlation values from a pixel by pixel analysis for OPP-mitochondrial colocalization relative to mean mitochondrial size, demonstrating a significant and positive correlation between increasing OPP colocalization and mitochondrial size (Regression line and 95% confidence intervals are plotted, data points from n>30 randomly imaged axons, selected from 3 replicate RGC preps; Pearson r was significant,*p < 0.05).

However, because of the potential variability with labeling efficiency, either from injection site differences, variations in injection volumes, and/or kinetics of intravitreal injection dispersion, we followed up in vivo experiments with in vitro approaches where EU and OPP labeling can be better controlled and quantified. For in vitro approaches, RGCs were isolated to 99% purity by immunopanning from early postnatal mouse retinas and seeded at low density, allowing for the visualization of individual cells and neurites. Then, cells were virally transduced to fluorescently label mitochondria with a Cox8a targeted dsRED, cultured for 48hrs, treated with TTX for 2hrs, and finally pulsed for 15 min or 1hr with OPP or EU, respectively. The cells were treated with TTX for a shorter period of time than during in-vivo experiments to avoid significant changes in viability, which at early postnatal days has the potential to decrease cell survival, unlike the in vivo time points tested^58^. However, no detectable decrease in viability after 2hr of TTX incubation was detected (Figure 6B). In RGCs in vitro, we again found cells with intensely labeled peri-nuclear regions, but no discernable differences in new transcript or protein levels in the cell body regions (Figure 6C), similar to in vivo experiments. Interestingly, in OPP-treated cells, obvious puncta were also visible in axonal segments. These puncta appeared scattered throughout distal axon segments in axonal tips, with varying sizes and numbers (Figure 6D). To see if activity inhibition by TTX could influence the presence of these axon-localized OPP puncta, we quantified the abundance of OPP sites within axonal segments, and found that TTX-treated axons demonstrate significant decreases (∼70%) in the number of OPP puncta as compared to controls, similar to the effects of translational inhibitor cycloheximide (Figure 6E), suggesting that activity regulates axon-localized protein synthesis. Note that these changes in puncta number are not likely due to decreased transport of newly synthesized proteins from the cell body, as the rate of soluble protein transport is less than 0.1 μm/s (or 90 μm in 15 min) and would be an unlikely captured change within the assay window in distal RGC axons^59^, which are typically longer than 600μm at 48hrs of culture in these cells^60^.

Axon-localized protein translation has been previously described^31^, but to further examine axon localized OPP puncta we immunostained cells with an antibody against cytoplasmic ribosomal protein S3, an integral component of the 40s-ribosomal subunit translation initiation site^61^. In these cells, we found a significant level of co-localization of ribosomal protein S3 with OPP puncta and interestingly a large majority of these OPP puncta also colocalized with mitochondria (Figure 6F,G), suggesting that the detected axon-localized OPP puncta active local translation sites are often at or near mitochondria. Similar to our in vivo results, we further found that mitochondrial size decreased in TTX-treated RGCs (Figure 6H) and was increased in the presence of BDNF, which was dominant over the effect of TTX, with increases or decreases in mitochondrial size correlating with increases or decreases in OPP-to-mitochondria localization (.499 r) (Figure 6J). Of note, these puncta were not visible in EU-treated cells, which is either a reflection of the abundance of RNA within axons or an indication of detection limitations. Thus new protein synthesis is detected at mitochondria and together with mitochondrial network size shows a dependence on electrical activity in RGC axons.

### RNA binding proteins tether nuclear encoded mitochondrial mRNA to mitochondria

To investigate the potential for mitochondria to act as docking sites for nuclear-encoded mitochondrial mRNA and new protein translation and, we conducted proteomic and qRT-PCR experiments on mitochondria purified from optic nerves or retinas. Briefly, isolation of axonal mitochondria was performed by incubating homogenized whole optic nerve tissue with a magnetically conjugated Translocase Of Outer Mitochondrial Membrane 22 (TOM22) antibody. Then, homogenates were passed through magnetic columns to remove cytosolic contaminants, followed by extensive washing, elution, and the pelleting of mitochondria^62^ (Figure 7A). We used a number of approaches to validate this relatively novel purification protocol. First, isolated mitochondria were examined by SEM and TEM, which showed that mitochondrial membranes and cristae architecture were structurally maintained after isolation with dark puncta visible on the outside of mitochondria (Figure 7B), representing nanoparticle-bound TOM22 and confirming the integrity of outer mitochondrial membranes after isolation. On western blots, isolated mitochondria maintained proteins from all complexes in the electron transport chain (Figure 7C) and inner and outer membrane integrity proteins (Figure 7D) but no detectable cytoplasmic GAPDH (Figure 7E). Of note, supernatant fractions from TOM22-purified mitochondria, which would reflect TOM22-bound membranes from ruptured, non-intact mitochondria, had no detectable ETC or membrane integrity proteins (Figure 7C, D), suggesting that nearly all TOM22-purified mitochondria were intact and captured in the pellet fraction. Finally, TOM22-selected optic nerve mitochondria were isolated from Thy-1-CFP/COX8a mice and further purified to axon-specific mitochondria through a traditional fluorescence acquired cell sorting (FACS) machine. In all FACs assays, mitochondria from Thy-1-CFP/COX8a mice retained membrane integrity proteins, detected by western blot (Figure 7F), and CFP positivity, detected by mitotracker-CMXROS co-staining(Figure 7G). Furthermore, isolated CFP+ mitochondria maintained high membrane potentials and readily took up JC-1, forming distinct populations of red shifted J-aggregate-retaining mitochondria (Figure 7H) that lost polarization in response to a membrane potential uncoupler, carbonyl cyanide-*4*-(trifluoromethoxy) phenylhydrazone (FCCP) (Figure 7I, J). Overall, these data suggest that isolating optic nerve axon mitochondria via magnetic columns and FACS yields relatively pure and structurally intact mitochondria, with surface and internal proteins and functional polarization maintained throughout the procedure.

**Figure 7.**
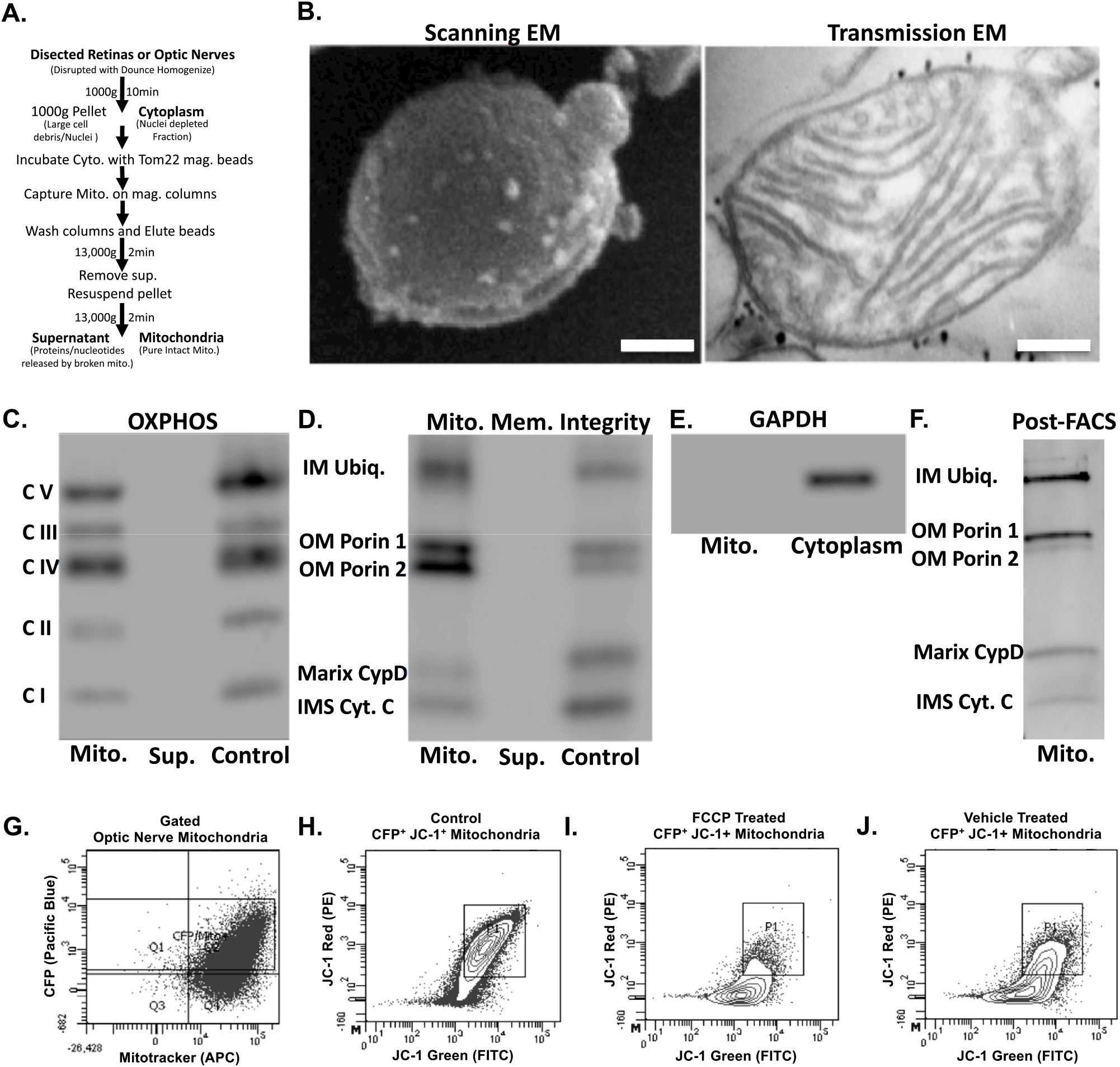
Purified mitochondria retain their protein content and membrane integrity. (A) Outline of mitochondrial isolation and subsequent assays. (B) TOM22-bound nanoparticles are visible, bound to the outer mitochondrial membrane in both SEM (white spots) and TEM (black dots). Scale bars 50 nm and 100 nm. (C-F) Western blot analyses of purified mitochondria and supernatants. Magnetically isolated mitochondria retain (C) OXPHOS subunits, as well as (D) outer membrane (OM), inner membrane (IM), and inner membrane space (IMS) proteins. (E) GAPDH is detectable in cytoplasmic but not mitochondrial isolate fractions. (F) FACS-sorted mitochondria retain both inner and outer membrane integrity proteins. (G) FACS-isolated mitochondria are intact and viable, retaining CFP and fluorescing with membrane potential-dependent mitotracker CMXROS. (H) Sorted CFP+ mitochondria demonstrate polarization-dependent fluorescence with JC-1, and (I) lose membrane potential with FCCP depolarization to a greater degree than (J) vehicle-treated controls.

We next examined these purified mitochondria for association with nuclear-encoded RNA binding proteins and translation-associated proteins from 3 young mouse optic nerves and also whole retinas by mass spectrometry, which yielded a total of 427 identifiable proteins after pooling all 6 samples. We cross-referenced these identified proteins with MitoCarta2.0, maintained by MIT’s Broad Institute, a database of thoroughly vetted proteins that co-purify with mitochondria^8,10^.

Using this approach, we were able to stratify our proteins into three groups of proteins based on evidence of mitochondrial localization (Figure 8A, B), 210 canonical mitochondrial proteins based on MitoCarta data from proteomics, computation, and microscopy analysis, 154 non-canonical mitochondrial proteins in the MitoCarta database that correlate with less pure mitochondrial fractions, and 63 proteins that are likely non-mitochondrial and do not show up in the MitoCarta database. Of course some of these “non-mitochondrial” proteins may still be mitochondrial as the MitoCarta data was not compiled from the visual system, where there could be uniquely-localized mitochondrial proteins.

**Figure 8.**
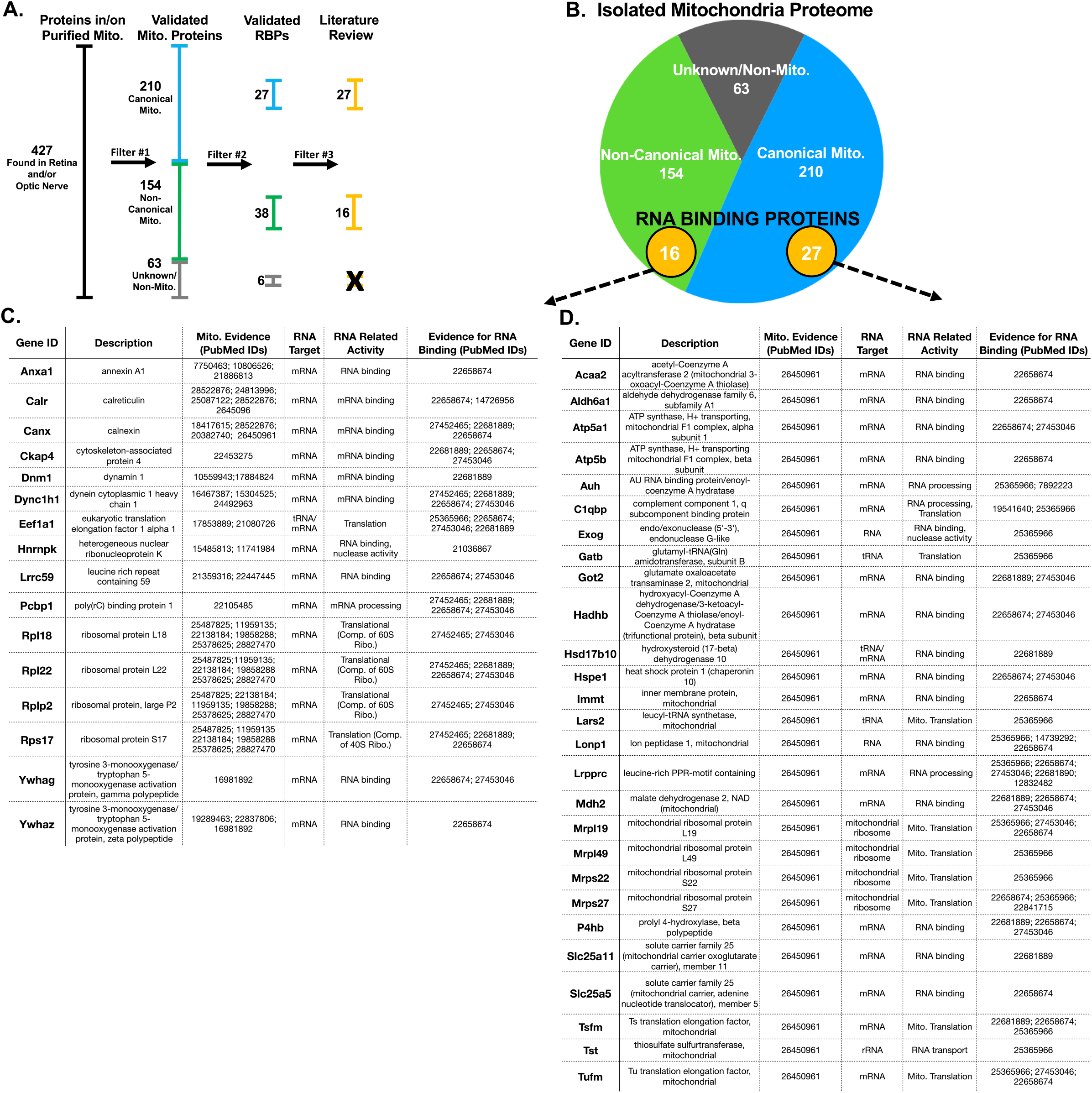
Proteomics mass spectrometry analysis reveals nuclear-encoded RNA binding proteins associated with purified mitochondria. (A) Filtering used to identify mitochondria-specific proteins and proteins with RNA binding properties.(B) Venn diagram of total protein hits sorted by annotation in the MitoCarta database. (C,D) Candidate mitochondria-associated RNA binding proteins with cited evidence (PubMed ID shown) for their functional RNA binding role and mitochondrial interaction (N=6 mitochondrial purifications).

After filtering for this external validation of mitochondrial association, we looked for the subset of proteins with predicted RNA binding potential by filtering proteins through David^63^, Panther^64^, and Uniport^65^ data bases using the gene ontology term RNA binding. Proteins that met these criteria were then cross-referencing against datasets from the RNA-protein interactome of cardiac cells^66^, HEK cells^67^, and HeLa cells^68^, as well as RNA binding protein databases AtTRACT and RBPDB^69-72^. This yielded the identification of 71 proteins with RNA binding properties, of which 43 had published evidence of direct interaction with mitochondria(Figure 8C, D). These included proteins with roles in translation, RNA processing, and RNA shuttling.

Since these data suggested that nuclear-encoded RNA binding proteins associate with mitochondria, we then asked whether there was also evidence for an association of nuclear-encoded mRNA on purified mitochondria. We purified mitochondria as above, extracted total RNA from pellets, and assayed for the presence of mRNA by qRT-PCR arrays. This identified a range of nuclear-encoded mitochondrial mRNAs, including genes that regulate mitochondrial dynamics, cell death, and energetics (Figure 9A,B). All detected mRNAs amplified in less than 30 cycles, lending confidence to the integrity of these measurements.

**Figure 9.**
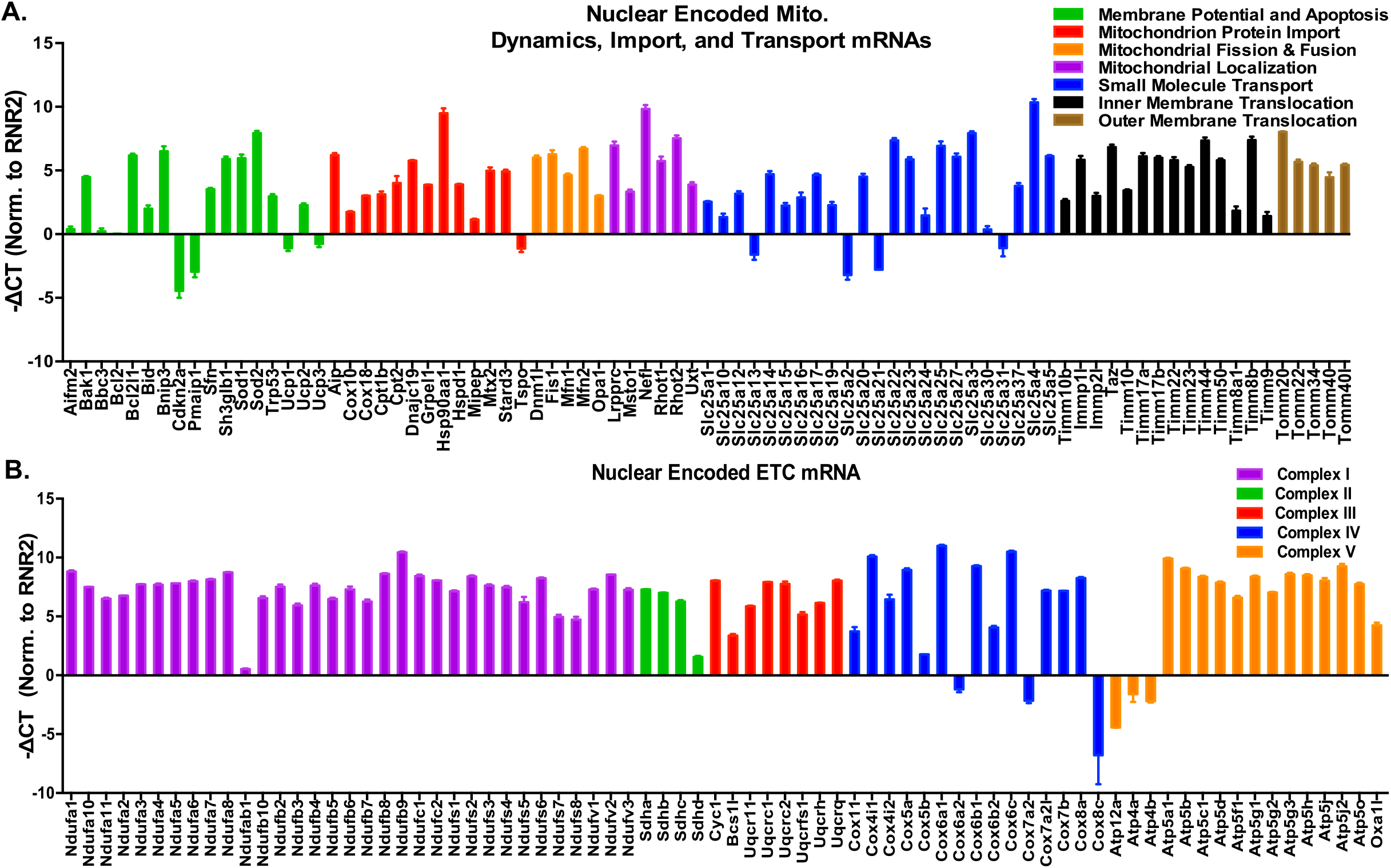
Isolated mitochondria bind nuclear-encoded RNAs associated with mitochondrial dynamics and energetics. (A) qRT-PCR array of mitochondrial dynamics and (B) energetics genes from purified mitochondria, normalized to RNR-2, a mitochondria-encoded ribosomal RNA(N=3 mito. purifications). All CT values were pulled from amplifications well below 30 cycles. The data is presented as toe inverse of die delta CT value, indicating more or less abundance of a particular nuclear encoded gene relative to RNR2. Genes are clustered by annotated functional roles in mitochondria.

To directly test the potential for mRNA tethering to outer mitochondrial membranes by RNA binding proteins, as implied by the OPP imaging experiments, mass spectrometry, and qRT-PCR array data, we treated mitochondrial isolates with Proteinase K to release surface proteins and associated mRNA, and then pelleted treated mitochondria to assay by qPCR for mRNAs released in the supernatant fractions (Figure 10A). As in Figure 8, to ensure that mRNAs detected in these assays were specific to mitochondria, we also verified the relative purity of mitochondrial fractions by western blotting for the presence of cytoplasmic contaminants, using actin an additional control (Figure 10B). We also took measures to verify that Proteinase K treatment did not release proteins from the interior of mitochondria, as shown by western blots against Complex III-Core Protein 2 and Complex V alpha subunit proteins (Figure 10C). In addition, there was no significant decrease in mitochondrial encoded mRNA ND4 in mitochondrial pellets and no significant increase in supernatant fractions (Figure 10D), providing strong evidence that Proteinase K did not interfere with inner mitochondrial protein or RNA. However, when purified mitochondria were tested for the release of nuclear-encoded mRNAs by Proteinase K, we found a significant number of mRNAs were released from mitochondrial pellets into the corresponding supernatant fraction, as compared to controls (Figure 10D). These mRNAs coded for proteins known to regulate mitochondrial dynamics, biogenesis, energetics, and RNA transport. We also found mRNA encoding cytoplasmic proteins GAPDH and actin bound to mitochondria, suggesting that bound mRNAs are not limited to those coding for mitochondrial-specific proteins. Interestingly, when we pre-treated in vivo with TTX at P11 and P13 and then assayed purified retinal mitochondrial transcripts from P15 mice, there were essentially no significant changes in mitochondrial localized mRNAs dependent on activity, although this could reflect an under sampling error as RGCs make up less than 1% of retinal cells. Overall, these data indicate that mitochondria bind nuclear-encoded mRNAs known to modulate mitochondrial size, number, and energetics, and that this process is mediated by RNA binding proteins present on mitochondrial membranes.

**Figure 10.**
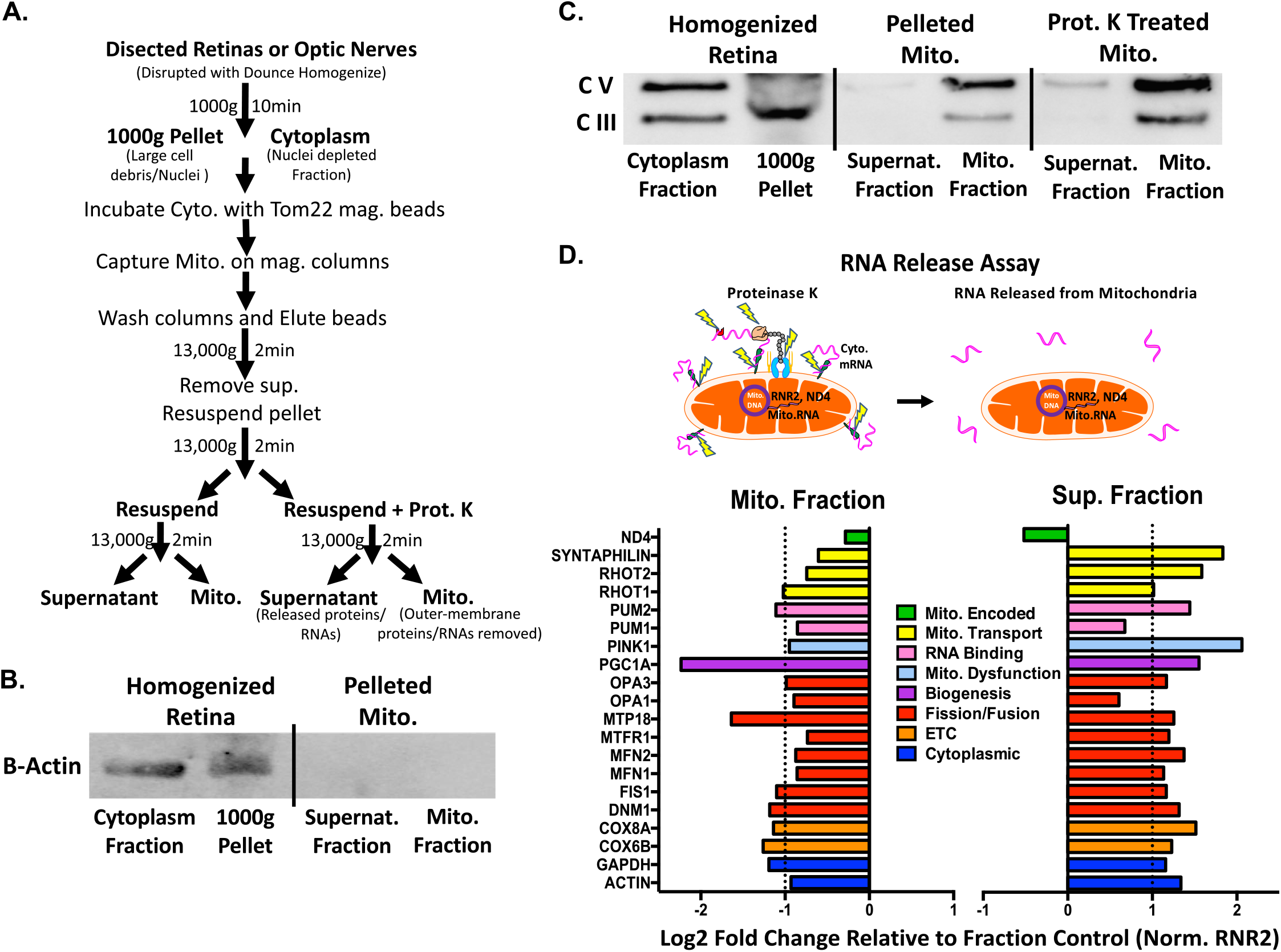
Purified mitochondria bind nuclear-encoded mRNAs through outer membrane-associated proteins. (A) Outline of mitochondrial isolation and subsequent assays. (B) β-actin is detected in homogenized retina but not in purified mitochondria. (C) Mitochondrial pellets treated with proteinase K retain inner matrix proteins, Complex Ill-Core Protein 2 (UQCRC2) and Complex V alpha subunit (ATP5A), confirming that proteinase K only strips off outer membrane-associated proteins. (D) Proteinase K-treated mitochondria release bound RNA into the supernatant fraction, as detected by qPCR of pelleted mitochondrial fraction and corresponding supernatant fractions. Data normalized to control mitochondrial fraction and RNR2, graphed as a log2 fold change to represent up and down regulation of RNA. Dotted line represents changes greater than 2-fold (n=3 replicates from a mito. purification).

## Discussion

Proper CNS neuron development and homeostasis depends critically on mitochondrial organization and function throughout distal axonal segments. Furthermore, mitochondria have to be capable of dynamically changing to meet intra-axonal demands distal to the cell body. As result, neurons and their distal segments have particularly demanding requirements for the active expression, trafficking, and assembly of nuclear-encoded-mitochondrial-macromolecules (proteins and mRNAs). Here we build upon the known mechanisms regulating rapid mitochondrial change in axons, and present new findings in which mitochondria size, number, and total area are regulated via activity and BDNF, and implicate a role for associated activity-regulated mitochondrial localized translation in regulating distal mitochondrial dynamics in CNS axons.

### Activity and BDNF regulate mitochondrial morphology and localization

Concomitant with eye opening, the visual system experiences increases in RGC electrical activity and BDNF expression, triggered by Ca^+2^ signaling and the activation of CREB-mediated transcription^52, 73-75^. Both activity and BDNF signaling play a pivotal role in axon development, including axon growth and presynaptic maturation, with mitochondrial dynamics and energetics having stereotyped roles in these developmental events (reviewed elsewhere^76^). Yet, the link between activity or BDNF signaling and changes in mitochondrial dynamics during CNS axon maturation in vivo had not been investigated. Here we show that RGC activity and downstream BDNF during eye opening is sufficient and necessary to increase mitochondrial size and number in RGCs’ optic nerve axons during development, and that activity also plays a similar, albeit more muted role, in regulating mitochondrial morphology in RGCs’ retinal axon segments. Whether this dependence is also observed during earlier periods of RGC development, e.g. when RGCs experience correlated waves of activity generated by amacrine cells before eye opening, or in other developing neurons, will be important questions to pursue.

In addition, we provide data supporting a new model in which activity and BDNF are modulating mitochondrial dynamics, biogenesis, and energetics in part through gene expression and local protein translation. Analysis of gene expression data pointed to the activation of transcriptional networks linked to PGC1-α and RICTOR by activity and BDNF during the period of eye opening, similar to findings in other neurons^77^. Furthermore, cellular energetics are linked to PGC1-α^78^ and RICTOR^79^ signaling, and our data reflected these findings^80^, and suggest that activity and BDNF also regulate basal respiration in RGCs, with BDNF increasing the maximum respiration capacity of RGCs regardless of activity inhibition. Thus, our data is indicative of a generalized increase in mitochondrial biogenesis activity in RGCs treated with BDNF (and vice versa with TTX treatment), a mechanism by which morphology and distribution along axons may be regulated. Overall, these data also support a pathway in which eye opening and subsequent increased neuronal firing signal to modulate the expression of mitochondrial related transcription, mitochondrial dynamics, and energetics. There may also be a direct protein-based signaling cascade triggered by activity and subsequent BDNF signaling onto mitochondria, since there is evidence that activity and BDNF increase respiration independent of nuclei in mitochondria-containing synaptosomal preparations^53,81,82^.

### Activity regulates a novel mechanism of mitochondrial localized translation

Axon segments can be up to a meter away from a neuron’s cell body and nucleus, presenting a challenge for signaling and subsequent renewal of proteins required for normal neuronal function. Axonal transport of nuclear-encoded proteins, in which nuclear proteins are translated in the perinuclear space and transported down axons at rates of up to 8 mm/day for soluble proteins (i.e. metabolic enzymes) or 100 and even 400 mm/day when associated with mitochondria or neuropeptide containing vesicles, respectively^59,83,84^, may not be quick enough to resupply distal axonal sites at times of rapid demand. In addition, nuclear-encoded mRNAs including some for mitochondrial proteins^24-26^ are transported to distal axon sites including in RGC axons in the mouse^27^, ready to be translated locally and on demand^23^, suggesting that mitochondria function is in part maintained by active axonal localized translation. Consistent with our findings, activity modulation and neurotrophic factors including BDNF regulate local translation in xenopus neurons^85,86^ and our extend this model showing that such translation occurs at mitochondria in an activity-dependent manner. The enrichment in mRNAs encoding mitochondrial proteins on or near mitochondria^28-32^ and the finding that ribosomes can directly bind to mitochondria membranes via TOM proteins and act in co-translation protein import in yeast^33-35^, together suggest a novel mechanism whereby increased neuronal firing and BDNF downstream signaling pathways directly regulate mitochondrial dynamics through modified local translation (Figure 12). This model will require additional molecular investigation in mammalian axons, such as the development of approaches to inhibit protein synthesis in specific subcellular compartments.

**Figure 11.**
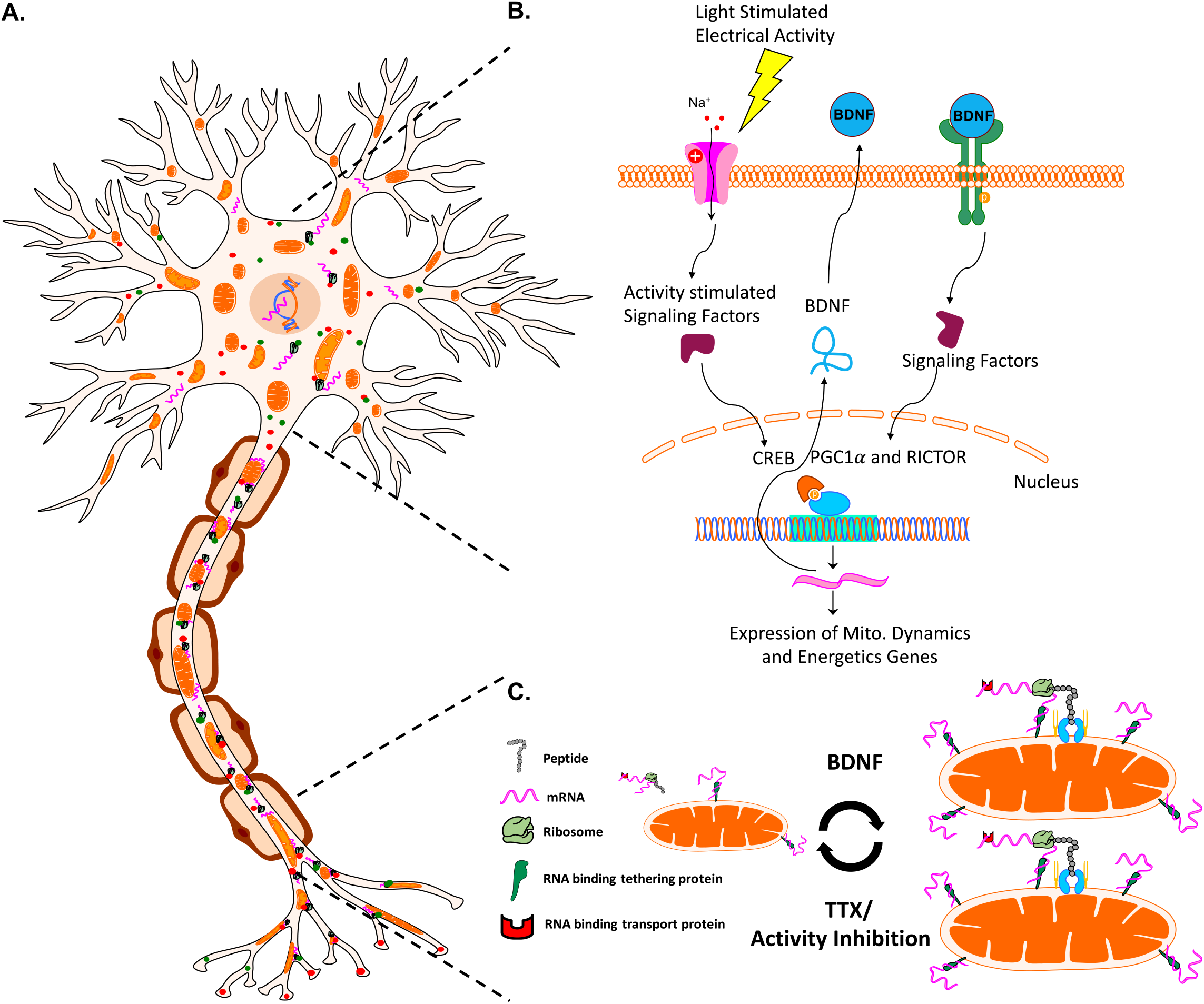
A model for activity- and BDNF-regulated mitochondrial size, number, and associated protein translation. (A) Neurons contain mitochondria, RNA, RNA binding and transport proteins, ribosomes, and newly synthesized proteins, throughout distal axon and dendrite compartments. (B) Electrical activity in RGCs, for example driven by light stimulation of retinal circuitry after eye opening, activates a signaling pathway that culminates in the activation of transcription factors such as CREB, and the expression of BDNF. BDNF signaling stimulates nuclear-encoded mitochondrial gene expression, coordinated by the activation of transcriptional regulators RICTOR and PGCl-α.(C) Neuronal activity and downstream BDNF signaling stimulates increases in mitochondrial size and number, reversed by activity inhibition. Changes in mitochondrial size also correlate with mitochondrial localized translation of nuclear-encoded transcripts

### Implications for aging and disease

These data move towards identifying mechanisms regulating mitochondrial organization and nuclear-encoded mitochondrial transcript localization in CNS axons during normal developmental, but raise questions to what degree similar mechanisms act in aging or neurodegenerative disease. Declining or defective mitochondrial function has been linked to many neurodegenerative diseases^87^. In humans and in mammalian animal models, defective axonal mRNA transport mechanisms have been implicated in the pathogenesis of neuropathies including spinal muscle atrophy, amyotrophic lateral sclerosis, and distal hereditary neuropathy^88-90^, and declining metabolic function is increasingly linked to reduced expression of mitochondrial transcripts^91,92^. Thus, understanding how the expression and local translation of nuclear mitochondrial transcripts are regulated and how these influence mitochondrial function may yield new approaches to treat dysfunction in the nervous system.

## Materials and Methods

### Animal use statement

Experiments conformed to the ARVO Statement for the Use of Animals in Ophthalmic and Vision Research and were approved by the Stanford University Biosafety Committee and the Institutional Animal Care and Use Committee. Strains used in these experiments included wild type CD-1, C57BL/6, and B6.Cg-Tg(Thy1-CFP/COX8A)S2Lich/J mice (The Jackson Laboratory).

### Cell Culture

RGCs were purified from male and female postnatal day 1/2 (P1/2) C57BL/6 mice (Charles River Laboratories) by immunopanning, and cultured on poly-D-lysine- (10 μg/mL; Sigma Aldrich, P-6407) and laminin-coated (2 μg/mL; Sigma, L-6274) cover glass bottom 96 well plates (Greiner Bio-One)^93,94^, in media with or without BDNF supplement as previously described^86^. Then, RGCs were treated with baculoviruses to label mitochondria (BacMam 2.0, Thermo Fisher Scientific, C10601) and 48 hrs later incubated with pharmacological agents, at stated concentrations and times. In RNA or protein labeling experiments RGCs were incubated with EU for 2 hrs or OPP for 15 min (according to the Click-iT Nascent RNA or Protein Synthesis Assay Kit from Thermo Fisher Scientific, C10327 or C10456). After all incubations cells were then fixated (4% PFA PBS), permeabilizated (0.5% Triton PBS), and in EU- or OPP-incubated cells, Click labeling reaction were performed. To identify ribosomes, cells were incubated with antibodies against ribosomal protein S3 (Cell Signaling Technology, D50G7) at 1:100 overnight at 4°C, and secondary Alexa-647 conjugated antibodies at 1:500 for 2hrs at room temperature. To label growth cones, cells were incubated with Alexa Fluor® 647 Phalloidin (Thermo Fisher Scientific, A22287) at 1:40 room temp. for 30 min prior to confocal imaging. In oxygen consumption experiments, RGCs were plated as described above, with or without BDNF, at 40k/well in 96 well plates designed for the Seahorse XF96 instrument (Agilent, 101085-004). After culturing for 24hrs, media was exchanged with Assay Media (Agilent, 102365-100) and FluxPak injectable ports were loaded with drugs as recommended by mitochondrial stress test kit (Agilent, 103015-100). TTX was loaded in the empty port A as the first injection, followed by Oligomycin, FCCP, and Rotenone/Antimycin A, respectively. After assay was complete oxygen consumption values were normalized to the number of Dapi positive cells per well.

### Imaging

B6.Cg-Tg(Thy1-CFP/COX8A)S2Lich/J mice (CFP/COX8a, Jackson labs) were euthanized and perfused with 4% PFA in PBS at P9, P13, P15, and P45. Perfused animals were then enucleated and the eyes and the optic nerves post-fixed in 4% PFA for 1-3 hours. Post fixed tissues were whole mounted on slides in Vecta-Shield mounting medium (Vector Labs, #H-1400) and imaged on a Zeiss LSM 710 confocal microscope. Compressed Z-stacks were analyzed by selecting nine random 25 × 50 μm sections and then measuring CFP expression with the ImageJ particle analyzer tool (National Institutes of Health). All images of cultured RGCs were collected on a Zeiss LSM 880 confocal system with a 40x/63x objective and using airyscan imaging mode, followed by airyscan processing using Zen software. Mitochondrial size, translation spot size, and colocalization analysis was conducted using Volocity Imaging Software (Perkin Elmer).

### Electron microscopy

Adult CD-1 mice under anesthesia were perfused with one half Karnovsky’s fixative; 2.5% glutaraldehyde and 2% paraformaldehyde (PFA) in 0.2M cacodylate buffer. Mice were euthanized and eyes with optic nerves were post fixed in half Karnovsky’s fixative. Tissues were placed in 2% glutaraldehyde overnight and then rinsed in 0.1M phosphate buffer with osmium tetraoxide. Osmicated tissues were rinsed in 0.15M phosphate buffer and dehydrated with graded concentrations of cold ethanol, ranging from 25 to 100%. Dehydrated tissues were rinsed with propylene oxide and embedded in Epon-Araldite with DMP-30 (All reagents were purchased from Electron Microscopy Sciences). Mitochondrial numbers were counted per axon area delineated by morphological features. Mitochondrial and axon boundaries were manually traced, and the areas were calculated using ImageJ analysis software (National Institutes of Health).

### Mitochondrial Purification

Whole retinas or optic nerves and tracts were quickly dissected from CO2 sacrificed mice, and homogenized using a dounce tissue grinder (Wheaton, 357538) with 20-30 strokes in mitochondrial isolation buffer (provided in the Mitochondria Isolation Kit, Miltenyi Biotec, 130-096-946) with protease inhibitors (Thermofisher Scientific, 78425). The homogenate was then spun at 1000g and the supernatant was removed for subsequent magnet based mitochondrial isolation according to Milteny Biotec’s Mitochondrial Isolation Kit. Isolated mitochondria were washed and re-pelleted three times to insure mitochondrial fractions were pure and intact, for all downstream experiments. In addition, all procedures were performed on ice or at 4°C, to preserve mitochondrial integrity.

### FACS analyses of mitochondria

CFP-expressing mitochondria were analyzed with forward and side scatter in a Becton Dickinson FACScan (Becton Dickinson, San Jose, CA). Data were acquired in list mode, evaluated with WinList software (Verity Software House). In some experiments, mitochondria were detected with anti-TOM20 antibodies (Abcam, ab78547) or mitotracker CMXROS (Thermofisher Scientific, M7512). To determine if mitochondria were intact and viable, FACS sorted mitochondria were equilibrated with the membrane potential–sensitive dye JC-1 (500 nM; 5,5’6,6-tetra-chloro-1,1,3,3-tetraethylbenzimidazol-carbocyanine iodide; Thermofisher Scientific, T3168) for 20 minutes with or without FCCP (10 μM; carbonylcyanide-P-trifluoromethoxyphenylhydrazone; Sigma Aldrich, C2920).

### Western Blots

To further evaluate the structural integrity and purity of isolated mitochondria, mitochondria were analyzed by western blot using the following antibodies. Antibodies against inner and outer mitochondrial membrane integrity proteins include; Outer Membrane - Porin (VDAC1), Inner Membrane - Ubiquinol Cytochrome C Reductase Core Protein I, Intermembrane Space - Cytochrome C and Complex Va, and Matrix Space-Cyclophilin 40, (Abcam-ab110414; ab14734, ab110252, ab110325, ab110273, and ab110324). Electron transport chain protein antibodies include; Complex I subunit (NDUFB8), Complex II-30kDa (SH3B), Complex III-Core Protein 2 (UQCRC2), Complex IV subunit I (MTCO1), and Complex V alpha subunit (ATP5A) (Abcam-ab1104; ab110242, ab14714, ab14745, ab14705, and ab14748). Cytoplasmic antibodies include; GAPDH and β-Actin (Cell Signaling Technology, 2118S and 8457). GFP/CFP antibodies (Thermofisher Scientific, A10262) were used as controls for the integrity of mitochondrial fractions collected from Thy1-CFP/COX8A mice.

### Proteomics

To detect mitochondrial associated proteins, mitochondrial purification was performed on 24 optic nerves and 6 retinas. Resulting mitochondrial pellets were solubilized with 5% rapigest in TNE buffer and then boiled for 5 min, followed by reduction in 1mM Tris(2-carboxyethyl)phosphine hydrochloride at 37°C for 30min. Then samples were alkylated in .5mM 2-iodoacetamide at 37°C for 30min, followed by trypsin digestions at 1:50 (enzyme:protein) overnight at 37°C and the addition of 250mM HCl at 37°C for 1hr. Samples were then centrifuged and peptides were extracted from the supernatant and desalted using Aspire RP30 desalting columns (Thermo Scientific)^95^. Trypsin-digested peptides were analyzed by LC-MS/MS^96^ on the TripleTOF™ 5600 hybrid mass spectrometer (ABSCIEX). MS/MS data were acquired in a data-dependent manner in which the MS1 data was acquired for 250 ms at m/z of 400 to 1250 Da and the MS/MS data was acquired from m/z of 50 to 2,000 Da. For Independent data acquisition (IDA) parameters MS1-TOF 250 milliseconds, followed by 50 MS2 events of 25 milliseconds each. The IDA criteria; over 200 counts threshold, charge state of plus 2-4 with 4 seconds exclusion window. Finally, the collected data were analyzed and normalized^97^ using MASCOT^®^ (Matrix Sciences) and Protein Pilot 4.0 (ABSCIEX) for peptide identifications normalized based on spectral abundance factors.

### RNA detection

To detect mitochondrial or nuclear-encoded mRNA transcripts, RNA was extracted from isolated RGCs or mitochondria from retina or optic nerve and tract using the RNeasy Plus Micro Kit (Qiagen, 74034). RNA isolates were then processed for RT^2^ Profiler™ PCR Arrays (Mouse Mitochondrial and Mitochondria Energy Metabolism, PAMM-087ZE and PAMM-008ZE). For mitochondrial RNA release assays, isolated mitochondria were resuspended in mitochondrial suspension buffer (provided in Mitochondria Isolation Kit) and incubated with Proteinase K (Thermofisher Scientific, 25530049) at 5ug/mL for 10 min. Mitochondria were then pelleted, and supernatant and mitochondrial pellets were separately processed for RNA purification and subsequent qPCR arrays. All qPCR data were acquired on QuantStudio 7 Flex Real-Time PCR System (Applied Biosystems, Thermofisher Scientific).

### Pharmacologic interventions

Mice were anesthetized with xylazine (10 mg/kg, IP) and ketamine (80 mg/kg, IP). Anesthetized mice were injected intravitreally (1-2 μl) with vehicle, Hank’s balanced salt solution (HBS, Invitrogen), BDNF (3μg/μl; Peprotech #450-02), tetrodotoxin (TTX; 3 μM, Sigma #T8024), or combined TTX (3 μM) and BDNF (3.3 μg/μl; Peprotech).

### Eyelid opening or suturing

For premature eyelid opening experiments, P10 mice were anesthetized as above and eyelids were gently pried open with forceps as described^98^. Eyes were then treated with sterile 2.5% hydroxypropyl methylcellulose (Goniosol, Akorn) every 12-18 hours to ensure eyes remained open and lubricated throughout the duration of the experiment. For extended eyelid closure experiments, P11 pups were anesthetized and two mattress sutures were placed along the eyelid margin to prevent eye opening as described ^99^. Animals were checked daily to ensure sutured eyes remained closed until euthanasia.

### Graphing and Statistics

Data presentation and statistical analysis was done in Prism (Graphpad). To compare quantitative variables, Student’s t-tests or ANOVA with post-hoc t-tests were done with a *p*-value < 0.05 indicating statistical significance.

## Acknowledgements

We gratefully acknowledge funding from the NIH R01-EY020913 (JLG), P30-EY014801 (University of Miami), P30-EY026877 (Stanford University), F31-NS087789 (AK), as well as unrestricted grants from Research to Prevent Blindness, Inc. We thank the following for their technical support; Peggy Bates for electron microscopy, George McNamara and Gabe Gaidosh for microscopy, Kristina Russano and Eleut Hernandez for animal husbandry, Oliver Umland for assistance in flow cytometry, and Majid Ghassemian for mass spectrometry support.

